# P5A-ATPase controls the ER translocation of Wnt in neuronal migration

**DOI:** 10.1101/2021.04.13.439568

**Authors:** Tingting Li, Xiaoyan Yang, Zhigang Feng, Wang Nie, Yan Zou

## Abstract

Wnt family are conserved secretory proteins required for developmental patterning and tissue homeostasis. The mechanisms underlying intracellular maturation and intercellular signal transduction of Wnt proteins have been extensively studied. However, how Wnt is targeted to the endoplasmic reticulum (ER) for processing and secretion remains elusive. Here we report that CATP-8/P5A-ATPase directs neuronal migration non-cell autonomously in *C. elegans* by regulating EGL-20/Wnt biogenesis. CATP-8 functions as a translocase to translocate EGL-20/Wnt nascent polypeptide into the ER by interacting with the hydrophobic core region of EGL-20 signal sequence. Such regulation of Wnt biogenesis by P5A-ATPase is conserved in human cells. These findings reveal physiological roles of P5A-ATPase in neural development and identify Wnt proteins as direct substrates of P5A-ATPase for ER translocation.

## Introduction

Wnt proteins comprise an evolutionarily conserved morphogen family which plays fundamental roles in development patterning and tissue homeostasis. Dysregulations of Wnt signaling are associated with a variety of developmental deficits and diseases (Clevers, 2006). Upon secreted from producing cells, Wnt proteins trigger a variety responses including cell fate specification, polarity, and migration to target cells. Secreted Wnt glycoprotein forms a globular secondary structure through intramolecular disulfide bonds among 24 highly conserved cysteines. Therefore, functional Wnt requires multi-layers of posttranslational modifications, such as acylation, glycosylation and sulfation, critical for correct folding and subsequent secretion (Willert and Nusse, 2012). The journey of Wnt processing, folding and trafficking starts in the ER. However, upon translation, how Wnt targets to the ER is largely unknown.

Secreted Wnt proteins influence neural connectivity by patterning neuronal migration, axon guidance, and synapse formation. The Q neuroblasts in *C. elegans* serve as an ideal model to study neuronal migration and Wnt signaling (Chai et al., 2018). The bilateral Q neuroblast on the left side (QL) and Q neuroblast on the right side (QR) are born between V4 and V5 seam cells in the posterior worm, on the left and right sides symmetrically (Sulston and Horvitz, 1977). Although QL and QR undergo similar cell divisions, they migrate towards opposite directions. QL migrates posteriorly, differentiating into PQR, PVM and SDQL neurons (Figure 1A and 1B), whereas QR migrates anteriorly, producing AQR, AVM and SDQR neurons. The migration of Q neuroblasts is activated by EGL-20/Wnt proteins secreted posteriorly (Coudreuse et al., 2006), a process mediated by the endocytosis of Wnt-transport factor MIG-14/Wls by Retromer complex as well as proper folding by protein disulfide isomerase PDI-1 (Coudreuse et al., 2006; Pan et al., 2008; Torpe et al., 2019; Yang et al., 2008). Binding of EGL-20/Wnt to Frizzled receptors LIN-17 and MIG-1 in the QL cell leads to activation of the canonical Wnt signaling with BAR-1/β-catenin stabilization and nuclear localization (Harris et al., 1996; Maloof et al., 1999; Sawa et al., 1996), inducing the transcription of target genes including *mab-5* (Harris et al., 1996; Salser and Kenyon, 1992; Whangbo and Kenyon, 1999). The transduced signal initiates cytoskeleton rearrangement through WAVE/WASP coordination and hippo kinase (Feng et al., 2017; Zhu et al., 2016). In addition, Q cell polarity is regulated by UNC-40/DCC, PTP-3 and MIG-21 for its exposure to EGL-20/Wnt (Honigberg and Kenyon, 2000; Sundararajan and Lundquist, 2012).

**Figure 1.**
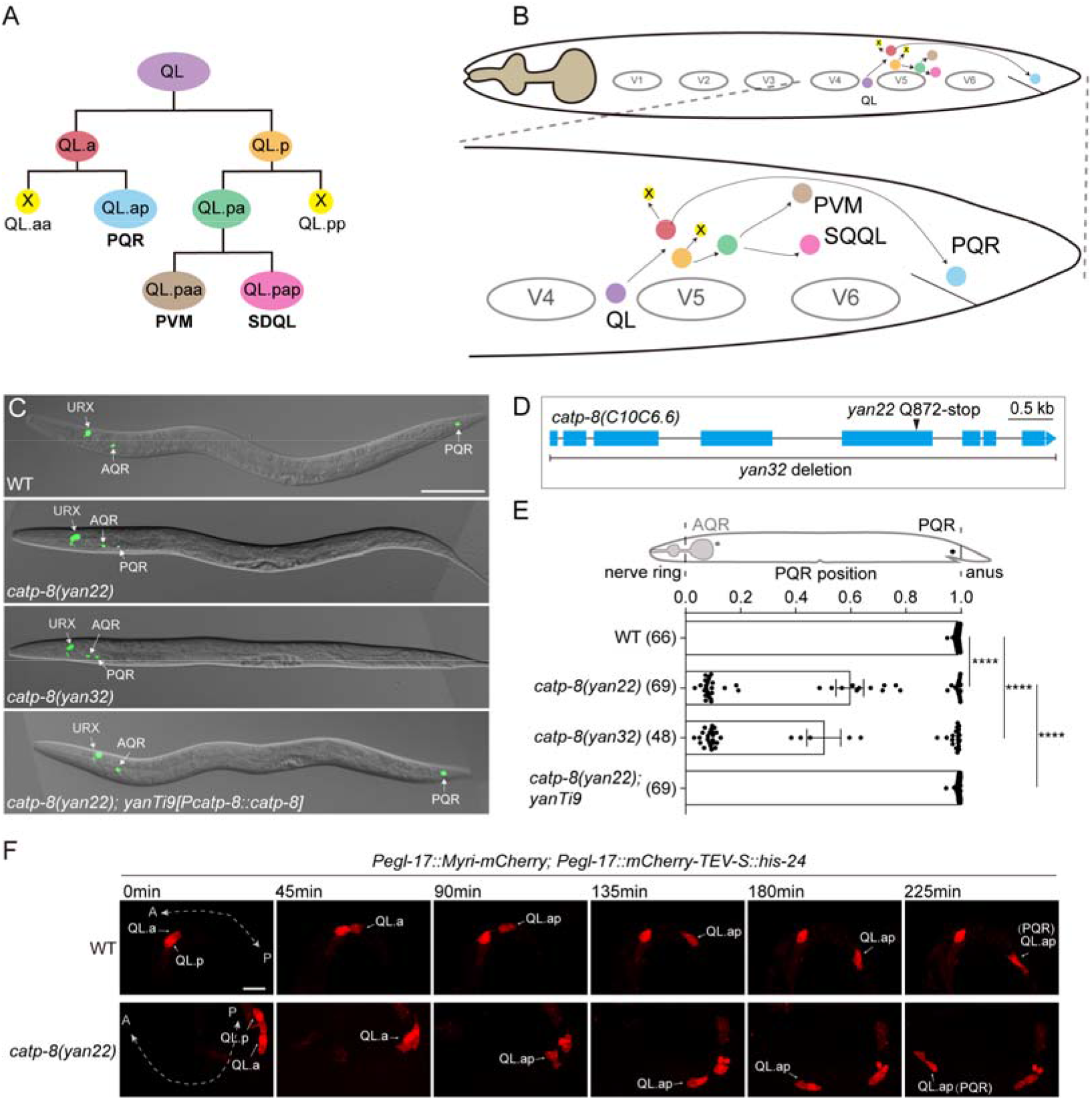
CATP-8/P5A-ATPase Is Required for QL Neuroblast Migration. (A) Schematic depicting QL neuroblast lineage. QL neuroblast undergoes three rounds of asymmetric divisions to yield two apoptotic cells (yellow circles with X) and three neurons PQR, PVM, SDQL. (B) Diagram showing cell divisions and migration of QL neuroblast. V1 through V6 depict the positions of six seam cells. Color code is the same as in (A). (C) Representative images of PQR positions in WT, *catp-8(yan22), catp-8(yan32)*, and *catp-8(yan22)* with a single copy transgene of *yanTi9 [Pcatp-8::catp-8]*. PQR neurons were visualized by a fluorescent marker *lqIs58[gcy-32::cfp]*. Scale bar, 100 μm. (D) Schematic of *catp-8* gene showing molecular lesions of *catp-8(yan22)* and *catp-8(yan32)* mutants. Arrowhead indicates the position of the point mutation in the *yan22* allele. (E) Quantifications of POR positions in (C). Data are presented as mean ± SEM. N numbers are shown in the brackets. ****, *P*<0.0001 (One-sided ANOVA with the Tukey correction). (F) Time-lapse imaging of QL descendant migrations at indicated time points after the first round of cell division. QL descendants were visualized by a fluorescent marker *rdvIs1[Pegl-17::Myr-mCherry; Pegl-17::mig-10::YFP; Pegl-17::mCherry-TEV-S::his-24]*. The dashed lines indicate worm positions with arrowheads pointing to A (Anterior) and P (Posterior) respectively. Scale bar, 10 μm.

P-type ATPases comprise a conserved transporter superfamily and are divided into five subfamilies: P1- through P5-ATPases. Typical P-type ATPases pump ions (P1- through P3-ATPases) or lipids (P4-ATPases) across cellular membranes with the energy of ATP hydrolysis (Palmgren and Nissen, 2011), while substrates of P5-ATPases have not been defined (Sorensen et al., 2015; Sorensen et al., 2019; Suzuki and Shimma, 1999). P5 subfamily consists of two subgroups: P5A-ATPase and P5B-ATPase. Recent studies provide insights into the substrates and gating mechanisms of P5-ATPases. Veen et al. demonstrate that ATP13A2, a human P5B-ATPase ortholog, transports lysosomal polyamine into the cytosol (van Veen et al., 2020). Next, we demonstrate that CATP-8/P5A-ATPase regulates dendrite branching by controlling the ER translocation of DMA-1 receptor in *C. elegans* (Feng et al., 2020). Another two papers report that P5A-ATPases safeguard ER integrity by removing mislocalized mitochondria tail-anchored (TA) proteins from the ER (McKenna et al., 2020; Qin et al., 2020). These findings suggest that P5-ATPases divert from other P-type subfamilies and have larger substrates in size like polyamines for P5B or polypeptides for P5A.

P5A-ATPase expression is abundant in mouse brain with the highest expression coinciding with the peak of neurogenesis (Weingarten et al., 2012). Moreover, human genetic analysis revealed that mutation in P5A-ATPase is associated with intellectual disability, attention deficit hyperactivity disorder (ADHD) and a host of developmental malformations and defects (Anazi et al., 2017). Despite the implication of P5A-ATPase in neural development, most studies on P5A-ATPase are carried out in yeast or cell culture. Only results from our previous work and Qin et al. have reported that dendrite branching in *C. elegans* requires P5A-ATPase to control the biogenesis of DMA-1 guidance receptor (Feng et al., 2020; Qin et al., 2020). However, DMA-1 is not very conservative to its mammalian ortholog. Therefore, physiological functions of P5A-ATPase in metazoans, particularly in the nervous system, remain to be elucidated.

Here, we report that the CATP-8/P5A-ATPase patterns neuronal migration in *C. elegans*. Our genetic and biochemical analyses reveal that that CATP-8 acts in Wnt-producing cells to control the ER translocation of EGL-20/Wnt. Interestingly, the hydrophobicity of core region in EGL-20 signal sequence determines the translocation dependency on P5A-ATPase. We further demonstrate that P5A-ATPase directly controls Wnt biogenesis in human cells in a conserved manner. Collectively, we identify Wnt as a P5A-ATPase substrate and is translocated into the ER for secretion to direct neuronal migration.

## Results

### CATP-8/P5A-ATPase Is Required for QL Neuroblast Migration

To understand the physiological role of P5A-ATPase in the nervous system, phenotypic investigation of *catp-8* mutants using different types of neuronal specific reporters was carried out. Whereas PQR neurons in wild-type (WT) animals are located next to the anus in the tail (Figure 1B), a majority of PQR neurons are mis-targeted and located near the anterior pharynx in the head of the worms carrying two *catp-8* null alleles (Figure 1C and 1E), *catp-8(yan22)* and *catp-8(yan32)* (Figure 1D). Transgenic *catp-8* expression driven by the endogenous promoter fully rescued the PQR migration defects in *catp-8* mutants (Figure 1C and 1E), suggesting that *catp-8* is required for PQR posterior localization.

As described above, PQR are differentiated from the QL neuroblast, accompanied by two other QL descendent cells, PVM and SDQL (Figure 1A). To explore whether migration of the whole QL lineage is affected by *catp-8* mutation, we also examined other QL lineage progenies: PVM and SDQL neurons. Similar to PQR neurons, both PVM and SDQL neurons migrate more anteriorly in *catp-8* mutants (Figure S1). Based on these results, we hypothesized that *catp-8* mutation impairs early migration of differentiated QL lineage progenies. To test this idea, we performed time-lapse imaging on the developing Q neuroblasts in live animals. As shown in Figure 1F, the first division descendants of QL neuroblast, QL.a and QL.p cells, were already oppositely migrated toward the anterior side in *catp-8* mutants, whereas their counterparts in WT animals protrude toward posteriorly. Taken together, our results suggest that *catp-8* is required to direct QL posterior migration.

### CATP-8/P5A-ATPase Acts in the Canonical Wnt Pathway to Control QL Migration

The asymmetric migration of QL and QR neuroblasts are determined by the intrinsic activation of the Hox transcription factor *mab-5*. The *mab-5*-expressing QL descendants migrate posteriorly, and QR progenies migrate anteriorly in the absence of *mab-5* (Salser and Kenyon, 1992). To uncover the genetic program *catp-8* mediating neuronal migration, we first tested the epistatic interaction between *mab-5* and *catp-8*. We found that *mab-5(lf);catp-8(yan22)* double mutants displayed similar QL migration defects to *mab-5(lf)* single mutants, while *mab-5(gf)* completely suppressed QL migration defects caused by *catp-8(yan22)* (Figure S2A), suggesting that *catp-8* acts upstream of *mab-5*. Next, we assessed whether *mab-5* is properly expressed in *catp-8* mutants using a *mab-5::gfp* reporter. Interestingly, *mab-5* expression was not detected in anteriorly-localized PQR cells in *catp-8* mutants, while posteriorly-localized PQR cells in *catp-8* mutants still expressed *mab-5* (Figure S2B). Therefore, defective PQR migration in *catp-8* mutants is likely due to the absence of *mab-5* expression.

The *mab-5*-dependent QL migration is activated by the canonical Wnt pathway (Harris et al., 1996). Signals relayed by the Wnt ligand EGL-20 are perceived by Frizzled receptors LIN-17 and MIG-1 in the QL cell, then transduced by MIG-5/Dishevelled to release BAR-1/β-catenin from PRY-1/Axin inhibition. Stabilized β-catenin is then translocated to the nucleus where it initiates the expression of many genes including *mab-5* (Figure 2A). To investigate whether *catp-8* acts through the canonical Wnt pathway to control QL migration, genetic interaction between *catp-8* and key components in the Wnt pathway was tested. We found that *catp-8* mutation enhanced the QL migration defect in Frizzled receptor mutant *lin-17* or *mig-1*, yet did not affect the QL migration in presence of *mig-5*/Dishevelled or *bar-1*/β-catemn mutation (Figure S2C). Notably, mutation of *pry-1*/Axin, the negative-acting factor in the Wnt pathway, suppressed the defective anterior displacement of QL progenies in *catp-8* mutants (Figure S2C). Taken together, these results suggest that *catp-8* acts upstream of *pry-1* in the canonical Wnt pathway to regulate QL migration.

**Figure 2.**
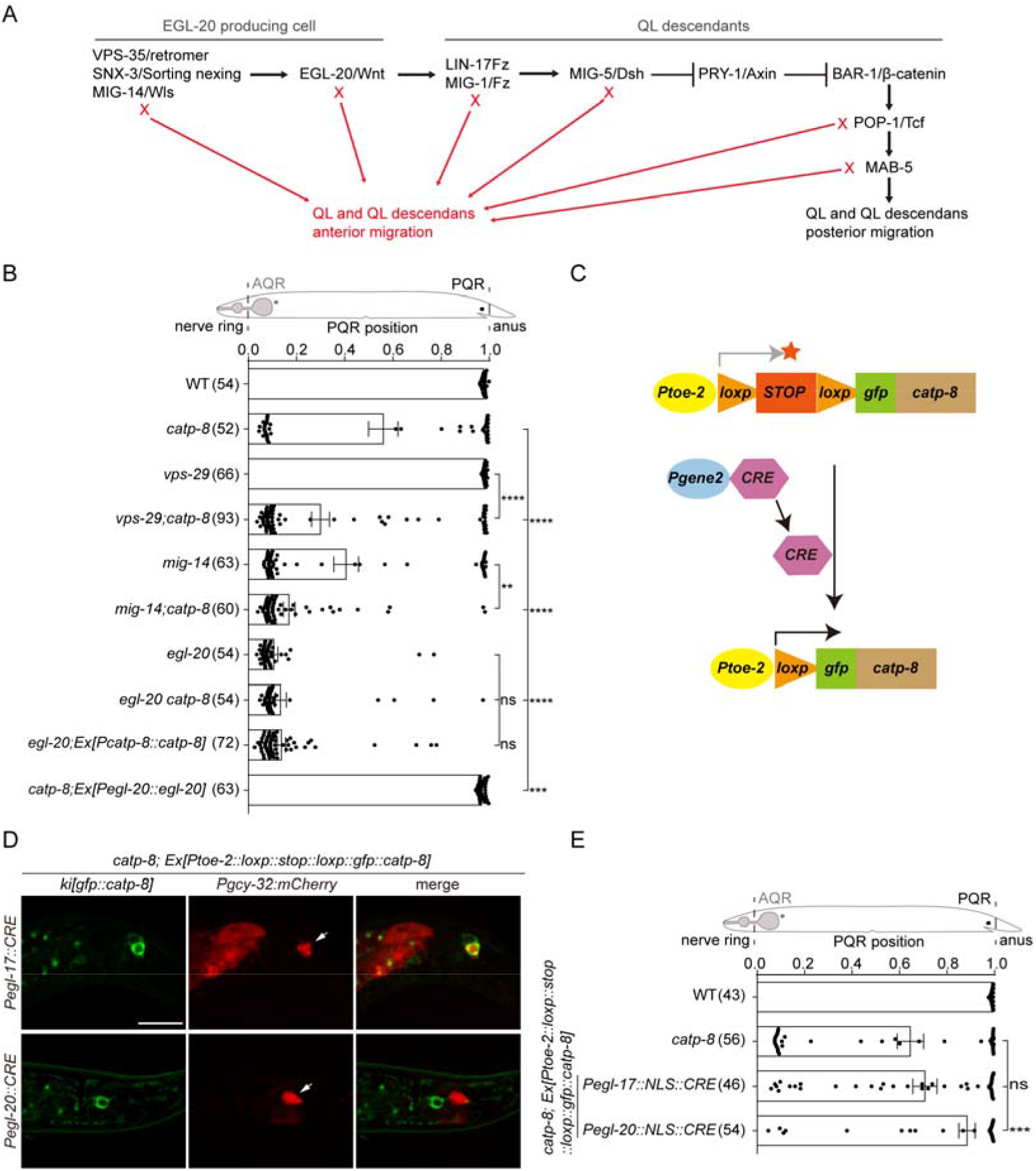
CATP-8/P5A-ATPase Acts in the Wnt Producing Cells to Control PQR Migration Non-Cell-Autonomously. (A) Schematic diagram of the canonic Wnt/β-catenin pathway required for QL migration, modified from (Eisenmann, 2005). (B) Quantifications of PQR positions in indicated genotypes, showing that *catp-8* acts upstream of *egl-20*. Data are presented as mean ± SEM. N numbers are shown in the brackets. **, *P*<0.01; ***, *P*<0.001; ****, *P*<0.0001; ns, not significant (One-sided ANOVA with the Tukey correction). (C) Diagram depicting the constructs of the Cre-dependent binary expression system used in (D). *Pgene2* refers to either *egl-17* promoter or *egl-20* promoter. (D) Representative images of *catp-8* expressions (Green) by the Cre-dependent binary expression system. PQR neurons were visualized by a *Pgcy-32::mcherry* transgene. Scale bar, 10 μm. (E) Quantifications of defective PQR migration in indicated genotypes, showing that overexpressing *catp-8* in Wnt-producing cells but not in PQR rescues PQR migration defects in *catp-8* mutants. Data are presented as mean ± SEM. N numbers are shown in the brackets. ***, *P*<0.001; ns, not significant (One-sided ANOVA with the Tukey correction).

To determine the role of *catp-8* in the canonical Wnt pathway, we further explored whether *catp-8* genetically interacts with known factors involved in EGL-20/Wnt biogenesis. As shown in Figure 2B, *catp-8* enhanced defective QL migration in anterior positions in *mig-14*/Wls and *vps-29*/Retromer but not *egl-20*/Wnt mutants, further supporting the notion that *catp-8* and *egl-20* act in the same pathway. Moreover, *egl-20* overexpression fully rescued QL migration defects in *catp-8* mutants (Figure 2B), further suggesting that *catp-8* functions upstream of *egl-20*/Wnt.

### CATP-8/P5A-ATPase Acts in Wnt-Producing Cells to Control Neuronal Migration Non-Cell-Autonomously

On the basis of our genetic results, we propose that *catp-8* acts non-cell-autonomously in the Wnt-producing cells to control QL migration. Using an N-terminal *gfp* knock-in strain *catp-8(yan27 ki[gfp::catp-8])*, we found that *catp-8* expressed ubiquitously in both PQR neurons and the surrounding hypodermis secreting Wnt (Figure S2D). Next, *catp-8* expression was driven under the *egl-20* or *egl-17* promoter in Wnt-producing cells or in QL descendants, respectively. To our surprise, *catp-8* overexpression driven by the *egl-20* promoter fully rescued the QL migration defects in *catp-8*, while *catp-8* overexpression under the *egl-17* promoter only partially rescued the defects (Figure S2E). *catp-8* from another Wnt-producing cell specific promoter, *Pspon-1* (Josephson et al., 2016), rescued the defects similarly as the *egl-20* promoter (Figure S2E). In addition, our analysis indicated that *egl-17* promoter drives a weak but broader expression in PQR neighboring cells (Figure S2D), attributing to the inconclusive and partial *catp-8* rescue results. To acquire exclusive expression of *catp-8* in QL descendants, a binary expression system combining *Pegl-17::Cre* and *Ptoe-2::loxp::stop::loxp::gfp::catp-8* was established (Figure 2C). The overlapping expression pattern of *toe-2* (Gurling et al., 2014) and *egl-17* promoters restricts the *catp-8* expression in QL descendants (Figure 2D). Interestingly, *catp-8* expression under the control of this system did not rescue the QL migration defects caused by *catp-8* (Figure 2E), indicating that *catp-8* expression in QL lineage is not sufficient. Altogether, our results suggest a non-cell-autonomous role of *catp-8* in QL migration.

### CATP-8/P5A-ATPase Regulates ER Translocation of EGL-20/Wnt through EGL-20 Signal Sequence

Since *catp-8* acts upstream of *egl-20*/Wnt in the EGL-20 producing cells, we next asked whether *catp-8* regulates *egl-20*/Wnt expression. Using an integrated *Pegl-20::egl-20::GFP* reporter, we found that *catp-8* depletion significantly reduced the EGL-20::GFP level, while re-expression of *catp-8* restored the reduction in EGL-20::GFP intensity (Figure 3A). Western blot analysis also revealed a reduction in EGL-20::GFP levels in *catp-8* mutants (Figure 3D), whereas *egl-20* mRNA levels measured by real-time RT-PCR were similar in both WT and *catp-8* mutants (Figure 3E). Based on these results, we conclude that *catp-8* regulates EGL-20 protein levels.

**Figure 3.**
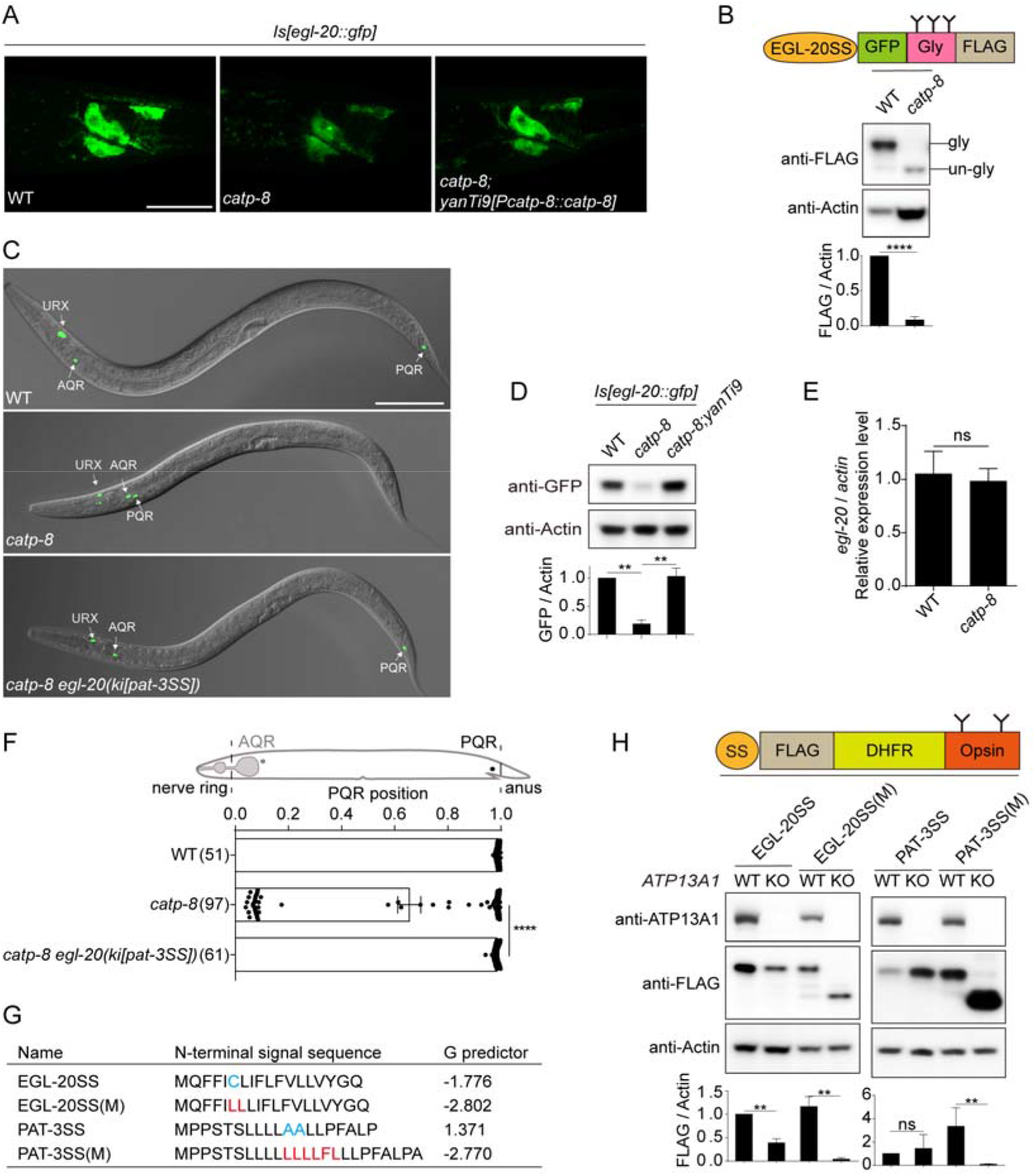
CATP-8/P5A-ATPase Controls ER translocation of EGL-20/Wnt. (A) Representative images of *egl-20::GFP* expression in WT, *catp-8(yan22)*, and *catp-8(yan22)* with a single copy transgene of *yanTi9 [Pcatp-8::catp-8]*. Scale bar, 20 μm. (B) Western blot of EGL-20SS-GFP-3*GLY-FLAG in WT and *catp-8(yan22)*. Bars indicate quantifications of the glycosylated proteins. Data are presented as mean ± SEM. ****, *P*<0.0001 (Student’s *t* test). (C) Representative images of PQR positions in WT, *catp-8(yan22)*, and *catp-8(yan22)* with a PAT-3SS knock-in to replace the endogenous EGL-20SS. PQR neurons were visualized by a fluorescent marker *lqIs58[gcy-32::cfp]*. Scale bar, 100 μm. (D) Western blot showing EGL-20::GFP protein levels in (A). Bars indicate quantifications of EGL-20::GFP and are presented as mean ± SEM. *, *P*<0.05 (Tukey’s multiple comparisons test). (E) Real-time RT-PCR showing mRNA abundance of *egl-20* in WT and *catp-8(yan22)* mutants. Data are presented as mean ± SEM. n.s., not significant (Student’s *t* test). (F) Quantifications of PQR positions in (E). Data are presented as mean ± SEM. N numbers are shown in the brackets. ****, *P*<0.0001 (One-sided ANOVA with the Tukey correction). (G) Signal sequences of EGL-20, PAT-33, and their mutants with higher hydrophobicity. The hydrophobicity was calculated by the Δ*G* prediction server v1.0 (https://dgpred.cbr.su.se/index.php?p=TMpred). (H) Western blot showing translocation efficiency of signal sequences in (G), assessed by the glycosylated band over Actin. DHFR is to generate a protein at suitable size. Opsin tag is a glycosylation reporter for protein entry into the ER (Judith Buentzel, 2017). Data are presented as mean ± SEM. **, *P*<0.01; ns, not significant (Student’s *t* test).

Previous study showed that CATP-8 is required for the biogenesis of DMA-1 receptor for dendrite branching in a signal sequence dependent manner (Feng et al., 2020). Upon translation, Wnt proteins are translocated into the ER and intracellularly processed and sorted for secretion (Willert and Nusse, 2012). We wondered whether CATP-8 controls the ER translocation of EGL-20/Wnt through EGL-20 signal sequence (EGL-20SS). To test this hypothesis, an engineered protein EGL-20SS::GFP::3*GLY::FLAG was constructed. If this engineered protein translocate into the ER, the three glycosylation sites (3*Gly) would become glycosylated and generate a larger protein than its predicted molecular weight. As expected, this EGL-20SS guided protein entered to the ER in WT animals but failed in *catp-8* mutants (Figure 3B), suggesting that its translocation is dependent on CATP-8. To confirm the requirement of EGL-20SS for CATP-8 regulated EGL-20 biogenesis, CRISPR-Cas9 approach was taken to replace the endogenous EGL-20SS with a previously described CATP-8 independent signal sequence, PAT-3SS (Feng et al., 2020). Indeed, PAT-3SS knock-in at the *egl-20* locus rescued QL migration in *catp-8* mutants (Figure 3C and 3F). Collectively, our data demonstrate that CATP-8 controls EGL-20/Wnt biogenesis through the recognition of specific signal sequence.

Signal sequences generally comprise characteristic tripartite architecture: a positive-charged N-terminal region, a hydrophobic core region and a C-terminal cleavage site by signal peptidases (Owji et al., 2018). Since both EGL-20SS and PAT-3SS only contain non-charged residues (Figure 3G), we wonder whether hydrophobicity is the key feature to confer CATP-8 dependence. We mutated EGL-20SS and PAT-3SS by increasing their hydrophobicity in the core region (Figure 3G), and then assessed their translocation efficiency in *ATP13A1*/P5A-ATPase knock out (KO) HEK293FT cells. We found that EGL-20SS(M) displayed higher dependence on ATP13A1 than EGL-20SS. Intriguingly, PAT-3SS(M) of increased hydrophobicity depended on ATP13A1 almost completely (Figure 3H). Taken together, these results suggest that P5A-ATPase recognizes signal sequences via the high hydrophobicity in the core region.

### P5A ATPases Control Wnt Biogenesis by Interacting with Wnt Signal Sequence in a Conserved Fashion

Since both P5A ATPase and Wnt proteins are present in higher organisms, we next asked whether the regulation of EGL-20/Wnt biogenesis by CATP-8 in *C. elegans* is evolutionarily conserved in mammals. To test this, we transfected human WNT1 in both WT and *ATP13A1* KO cells and found that WNT1 protein levels were dramatically reduced in *ATP13A1* KO cells (Figure 4A). By assessing WNT1 protein levels in media, we confirmed that WNT1 secretion was also severely impaired by *ATP13A1* mutation (Figure 4A). Moreover, similar to the observation in *C. elegans*, WNT1 signal sequence (WNT1SS) is essential for *ATP13A1*-dependent WNT1 biogenesis (Figure 4B). These results reveal the functional conservation of P5A-ATPases in regulating Wnt biogenesis.

**Figure 4.**
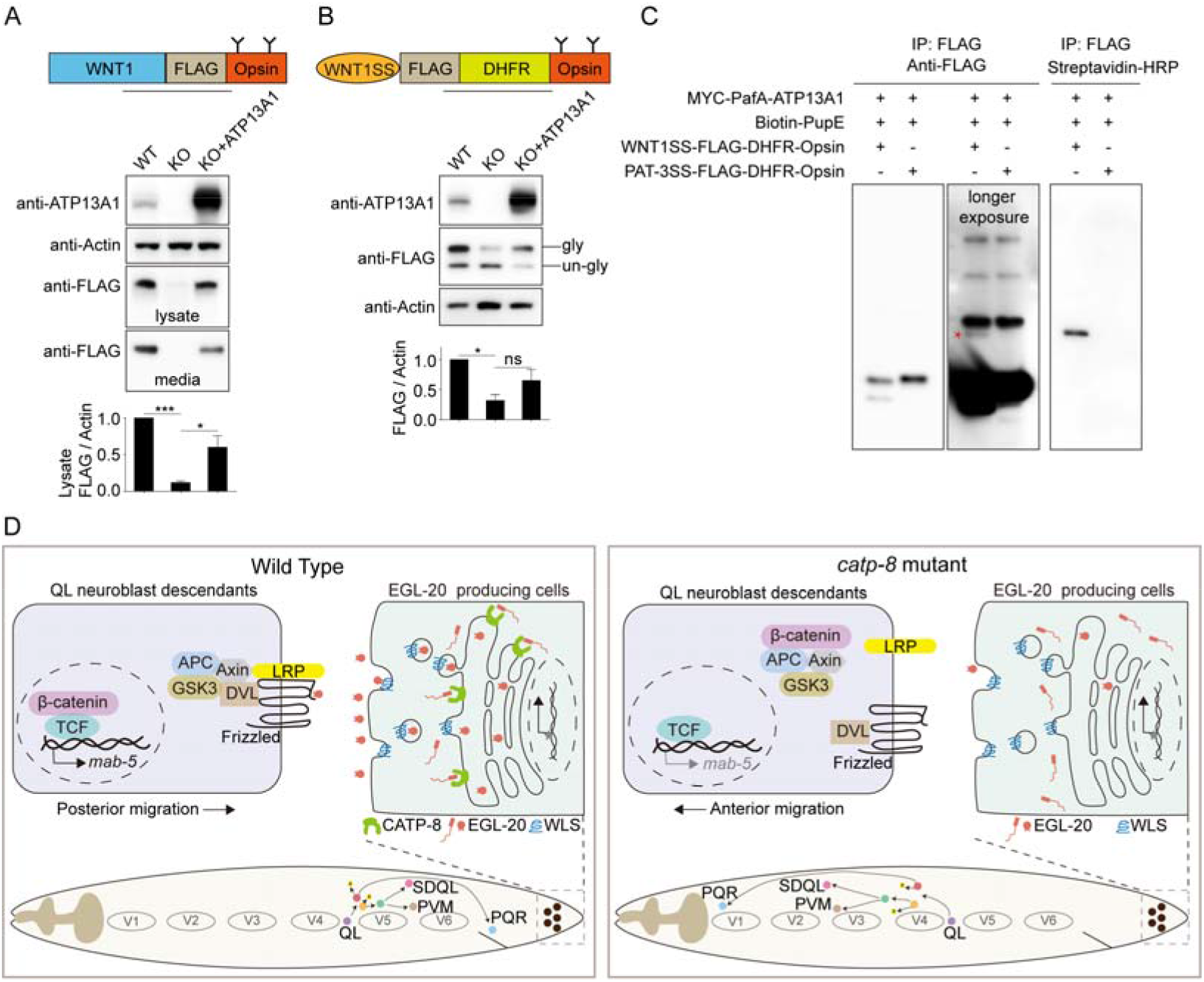
P5A-ATPases function conservatively to regulate Wnt biogenesis. (A) Western blot of transfected WNT1 in control, *ATP13A1* KO, and *ATP13A1* KO cells with *ATP13A1* overexpression. Data are presented as mean ± SEM. *, *P*<0.05; ***, *p*<0.001 (Tukey’s multiple comparisons test). (B) Western blot of transfected WNT1SS-FLAG-DHFR-Opsin in control, *ATP13A1* KO, and *ATP13A1* KO cells with *ATP13A1* overexpression. Data are presented as mean ± SEM. *, *P*<0.05; ns, not significant (Tukey’s multiple comparisons test). (C) Western blot showing proximal labeling of WNT1SS-FLAG-DHFR-Opsin by PafA-ATP13A1. Asterisk indicates Biotin-PupE labeled WNT1SS-FLAG-DHFR-Opsin. (D) Model for CATP-8/P5A-ATPase promoting ER translocation of EGL-20/Wnt through highly hydrophobic signal sequence of EGL-20.

We next sought to determine whether ATP13A1 mediates WNT1 translocation directly. As translocation is a very transient process, it is difficult to capture the interactions by co-immunoprecipitation. Thus, we employed the PUP-IT proximal labeling system (Liu et al., 2018) to detect whether there are physical interactions between ATP13A1 and WNT1. We transfected ATP13A1 fused with the proximity ligase PafA as a bait and either WNT1SS or PAT-3SS as a prey. If WNT1SS is the substrate of ATP13A1, PafA-ATP13A1 will catalyze the ligation of a small protein Biotin-PupE to Wnt1SS guided protein. Indeed, WNT1SS but not PAT-3SS guided protein was labeled with Biotin-PupE by PafA-ATP13A1 (Figure 4C), suggesting a direct and specific control of WNT1 translocation by ATP13A1.

## Discussion

In the present study, we identify CATP-8/P5A-ATPase as a key factor to direct neuronal migrations through regulating EGL-20/Wnt biogenesis. Our genetic and biochemical analyses demonstrate that CATP-8/P5A-ATPase controls the ER translocation of EGL-20/Wnt by specifically recognizing the hydrophobic core region in signal sequences (Figure 4D). Moreover, P5A-ATPase also regulates Wnt biogenesis in human cells, implying a conserved role of P5A-ATPase on Wnt-mediated developmental events.

Although numerous studies have focused on Wnt trafficking and signal transduction, it is still not clear how Wnt enters the ER for subsequent processing and secretion. Wnt family proteins harbor N-terminal signal sequences, and the signal sequence guiding Wnt ER translocation is thought to be similar to other secreted proteins. For those proteins, N-terminal signal sequences are recognized by Signal recognition particle (SRP) upon translation, and proteins are translocated into the ER through the translocation machinery Sec61 at the ER membrane. Our studies suggest that P5A-ATPase is a novel ER translocase based on its necessity of protein translocation and its physical interactions with signal sequences. P5A-ATPase emerges in the eukaryotes and exhibits distinct capacity for larger substrates like polypeptides (Feng et al., 2020; McKenna et al., 2020; Qin et al., 2020). Why translocation of proteins like Wnt requires P5A-ATPase is unclear. Interestingly, substrates of P5A-ATPase in metazoans, Wnt and DMA-1, are important for cell-cell communications and their biosynthesis is in high demand in relatively short developmental critical windows for a small group of cells. For example, EGL-20/Wnt is secreted from several cells in the worm tail and forms a gradient to pattern multiple early developmental events. DMA-1 is synthesized by two tiny PVD neurons and trafficked onto huge dendritic arborizations covering the whole worm body to perceive guidance cues from hypodermis (Dong et al., 2013; Liu and Shen, 2011). In these contexts, P5A-ATPase may function as a freeway to the ER for high demand proteins to be efficiently translocated, or to prevent the jam by heavy protein load to the essential translocon Sec61.

Intriguingly, different substrates of P5A-ATPase are likely to be transported in opposite directions. First, our studies demonstrate that P5A-ATPase directly controls the translocation from the cytosol to the ER of certain proteins with highly hydrophobic N-terminal signal sequences. Second, McKenna and Qin et al. show that P5A-ATPase functions as a dislocase to remove mistargeted mitochondria TA proteins from the ER to the cytosol (McKenna et al., 2020; Qin et al., 2020). Proteins with N-terminal signal sequences are thought to be generally less hydrophobic than TM and often have positive-charged N-termini, which might lead to their ER insertion in a wrong topology. It is proposed that P5A-ATPase might also remove these wrong topological proteins from the ER, providing additional opportunities for correct ER targeting (McKenna et al., 2020). However, our results define new features of signal sequences to confer P5A-ATPase dependence: high hydrophobicity and no-positive charged residues. Intriguingly, our data show that the vast majority of EGL-20 translocation depends on CATP-8, which suggests that CATP-8 is unlikely to function as a dislocase to remove wrongly targeting EGL-20. Otherwise, it seems inefficient and at high cost that most of EGL-20 proteins target to the ER in wrong topology by default and need to be removed by P5A-ATPase repeatedly until they target correctly. Therefore, P5A-ATPase is more likely to function as a translocase to directly translocate proteins into the ER. The mechanistic basis how P5A-ATPase recognizes different substrates and thus transports them in the opposite directions requires further study. The substrate specificity of P5A-ATPase might come from high hydrophobicity of signal sequences, as enhancement of hydrophobicity increases their translocation dependence on P5A-ATPase.

Directed neuronal migration is critical for the development of functional nervous system. Wnt proteins are known to play important roles in neuronal migration across species (Eisenmann, 2005; Yang et al., 2013). We identified Wnt proteins as direct substrates of P5A-ATPases in neuronal migration, underlying the mechanism of human ATP13A1 in neurodevelopmental disorders (Anazi et al., 2017). Considering ubiquitous expressions and pleiotropic phenotypes of P5A-ATPase (Anazi et al., 2017; Feng et al., 2020; Qin et al., 2020; Sorensen et al., 2015; Weingarten et al., 2012), more targets of P5A-ATPase in other cellular contexts remain to be identified for better understanding of the physiological roles of P5A-ATPase in metazoans.

## ACKNOWLEDGEMENTS

We thank Drs. Zhiyong Shao and Yingchuan Qi for reagents. We thank Drs. Margaret S. Ho, Yingchuan Qi, and Huanhu Zhu for instructive suggestions on the manuscript. The Molecular Imaging Core Facility (MICF) at ShanghaiTech University provided training on confocal microscopy. Some strains were provided by the CGC, which is funded by NIH Office of Research Infrastructure Programs (P40 OD010440). This study was supported by the National Natural Science Foundation of China (No.31571047) and the Start-up grant from ShanghaiTech University.

## AUTHOR CONTRIBUTIONS

Conceptualization, T.L., Z.F. and Y.Z.; Investigation, T.L., Z.F., X.Y., W.N. and Y.Z.; Writing – Original Draft, Y.Z. and T.L.; Writing – Review & Editing, Y.Z.; Funding Acquisition, Y.Z.; Supervision, Y.Z.

---

**Figure S1.**
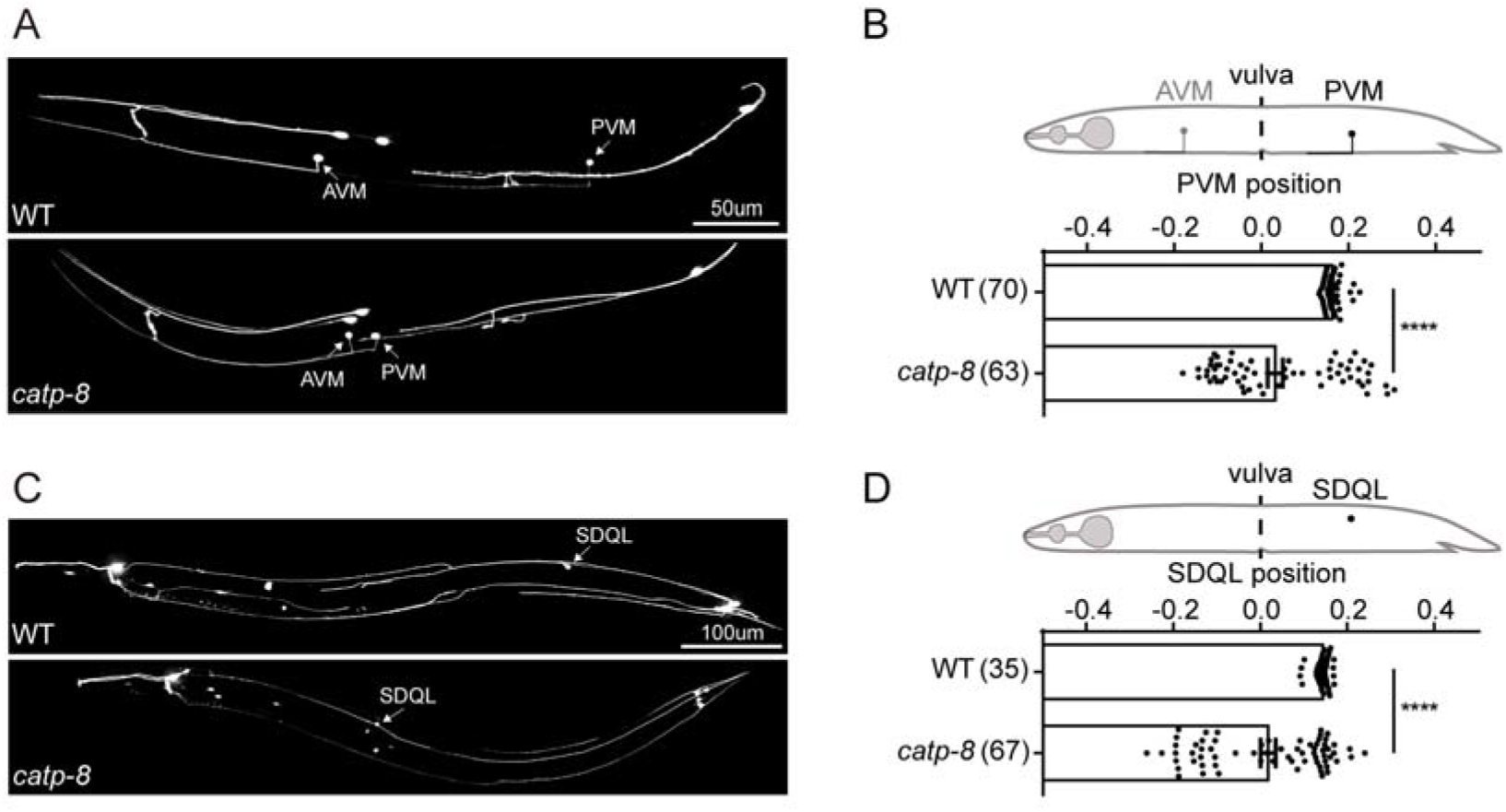
CATP-8/P5A-ATPase Is Required for PVM and SDQL Migration, Related to Figure 1. (A) Representative images of PVM positions in WT and *catp-8(yan22)*. PVM neurons were labeled by a fluorescent marker *zdIs5[Pmec-4::gfp]*. Scale bar, 50 μm. (B) Quantifications of PVM positions in (A). Data are presented as mean ± SEM. N numbers are shown in the brackets. ****, *P*<0.0001 (Student’s *t* test). (C) Representative images of SDQL positions in WT and *catp-8(yan22)*. SDQL neurons were labeled by a fluorescent marker *Ex[Pgcy-35::mcherry]*. Scale bar, 50 μm. (D) Quantifications of SDQL positions in (C). Data are presented as mean ± SEM. N numbers were shown in the brackets. ****, *P*<0.0001 (Student’s *t* test).

**Figure S2.**
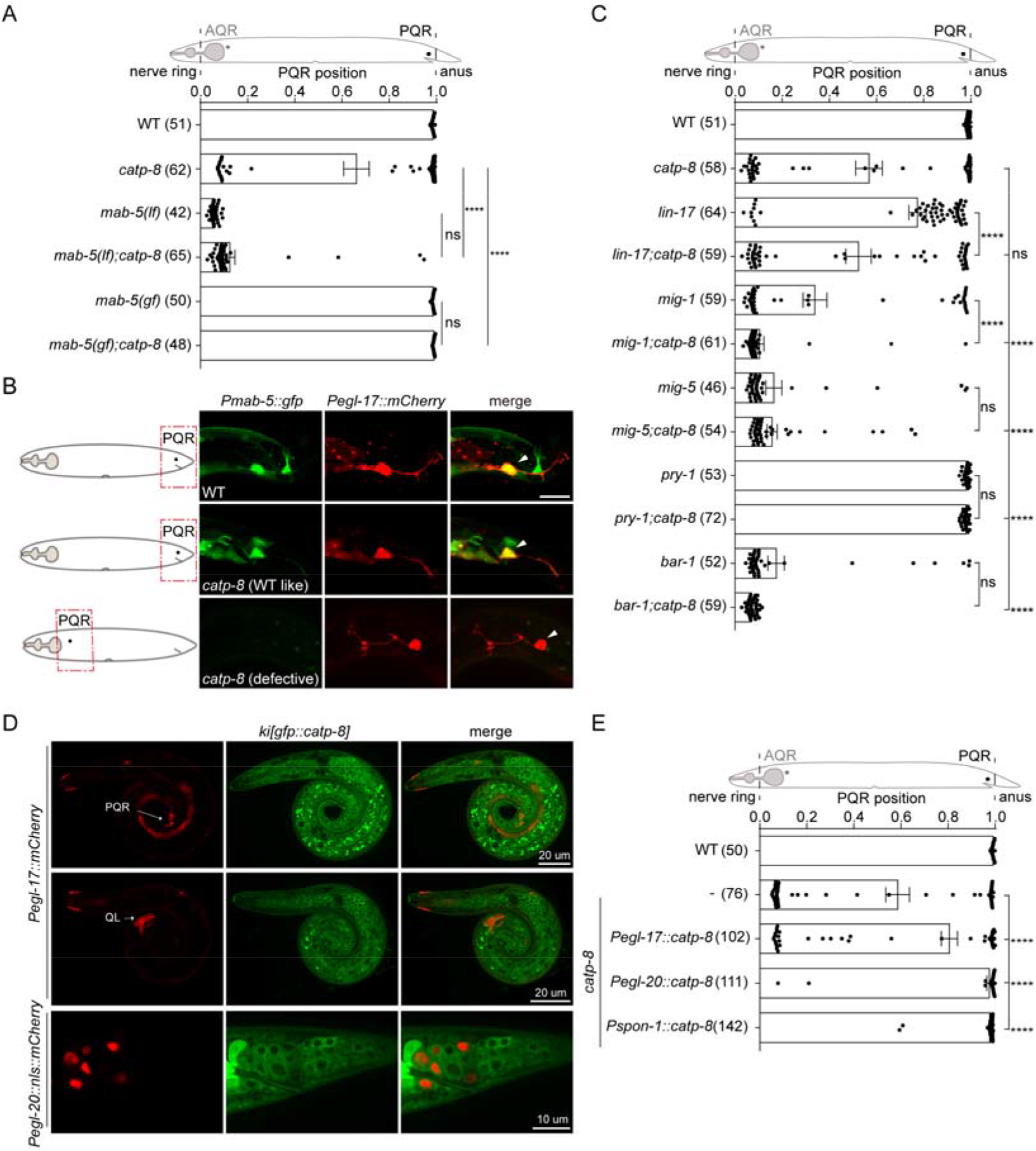
CATP-8/P5A-ATPase Acts in the Wnt Producing Cells to Control PQR Migration, Related to Figure 2. (A) Quantifications of PQR positions in indicated genotypes, showing that *catp-8* acts upstream of *mab-5*. Data are presented as mean ± SEM. N numbers are shown in the brackets. ****, *P*<0.0001; ns, not significant (One-sided ANOVA with the Tukey correction). (B) Representative images of *mab-5* expression shown by a *Pmab-5::gfp* transgene in WT and *catp-8* mutants. Diagram showing PQR positions in indicated genotypes. PQR neurons were visualized using a *Pegl-17::mcherry* fluorescence marker. Arrowheads point to PQR. Scale bar, 10 μm. (C) Quantifications of PQR positions in indicated genotypes, showing that *catp-8* acts upstream of the canonical Wnt pathway. Data are presented as mean ± SEM. N numbers are shown in the brackets. **, *P*<0.01; ***, *P*<0.001; ****, *P*<0.0001; ns, not significant (One-sided ANOVA with the Tukey correction). (D) Representative images showing CATP-8 expression in both Wnt producing cells and Q neuroblast. Green: CATP-8 expression shown by *ki[gfp::catp-8]*. Red: expression of *egl-17* promoter in PQR and the midline shown by *rdvIs1* (top), expression of *egl-17* promoter in QL shown by *rdvIs1* (middle), expression pattern of *egl-20* promoter shown by *Pegl-20::NLS-mCherry* (bottom). (E) Quantifications of PQR positions in indicated genotypes. Data are presented as mean ± SEM. N numbers are shown in the brackets. ****, *P*<0.0001 (One-sided ANOVA with the Tukey correction).

KEY RESOURCES TABLE

| REAGENT or RESOURCE | SOURCE | IDENTIFIER |
| --- | --- | --- |
| **Antibodies** |  |  |
| Mouse anti-FLAG | Sigma | Cat# F1804;<br>RRID:AB_262044 |
| Mouse anti-GFP | Abmart | Cat# M20004;<br>RRID:AB_2619674 |
| Rabbit anti-ATP13A1 | Proteintech | Cat# 16244-1-AP;<br>RRID:AB_10015282 |
| Donker anti-Rabbit | Jackson<br>Immunoresearch | Cat# 711-035-152;<br>RRID:AB_10015282 |
| Rabbit anti-Mouse | Sigma | Cat# A9044;<br>RRID:AB_258431 |
| Rabbit anti-b-Actin | Marine Biological<br>Laboratory | Cat# PM053-7;<br>RRID:AB_10697035 |
| Monoclonal ANTI-FLAG(R) M2,<br>clone M2 | Sigma | Cat# A2220 |
| **Bacterial and Virus Strains** |  |  |
| OP50 | CGC | http://www.cgc.edu/strain/OP50 |
| **Chemicals, Peptides, and Recombinant Proteins** |  |  |
| KOD-Plus-Neo | TOYOBO | Cat# KOD-401 |
| KOD One™ PCR Master Mix | TOYOBO | Cat# KMM-101 |
| Experimental Models: Cell Lines |  |  |
| HEK293FT | ATCC | RRID:CVCL_6911 |
| **Experimental Models: Organisms/Strains** |  |  |
| *C. elegans* strain, see Table S1 | This paper | N/A |
| **Oligonucleotides** |  |  |
| see Table S2 | This paper | N/A |
| **Recombinant DNA** |  |  |
| Plasmid: *pDD122* | Addgene | RRID:Addgene_47550 |
| Plasmid: *pDD162* | Addgene | RRID:Addgene_47549 |
| Plasmid: *Pgcy-32::mcherry* | This paper | N/A |
| Plasmid: *Pgcy-35::mcherry* | This paper | N/A |
| Plasmid: *Pcatp-8::catp-8* | This paper | N/A |
| Plasmid: *Pegl-17::catp-8* | This paper | N/A |
| Plasmid: *Pspon-1::catp-8* | This paper | N/A |
| Plasmid: *Pegl-20::catp-8* | This paper | N/A |
| Plasmid: *Pegl-20::nls::mcherry* | This paper | N/A |
| Plasmid:<br>*Ptoe-2::loxp::stop::loxp::gfp::catp-8* | This paper | N/A |
| Plasmid: *Pegl-20::nls::CRE* | This paper | N/A |
| Plasmid: *Pegl-17::nls::CRE* | This paper | N/A |
| Plasmid: *Pegl-20::egl-20* | This paper | N/A |
| Plasmid:<br>*Prpl-28::egl-20SS::gfp::Gly::FLAG* | This paper | N/A |
| Plasmid:<br>*pEGFP.C1-egl-20ss::FLAG::DHF*<br>*R::Opsin* | This paper | N/A |
| Plasmid:<br>*pEGFP.C1-egl-20ss(M)::FLAG::D*<br>*HFR::Opsin* | This paper | N/A |
| Plasmid: *pEGFP.C1-*<br>*pEGFP.C1-pat-3ss::FLAG::DHFR*<br>*::Opsin* | This paper | N/A |
| Plasmid: *pEGFP.C1-*<br>*pEGFP.C1-pat-3ss(M)::FLAG::D*<br>*HFR::Opsin* | This paper | N/A |
| Plasmid:<br>*pEGFP.C1-WNT1SS::FLAG::DHF*<br>*R::Opsin* | This paper | N/A |
| Plasmid:<br>*pEGFP.C1-WNT1::FLAG::Opsin* | This paper | N/A |
| Plasmid:<br>*pEGFP.C1-MYC::PafA::ATP13A1* | This paper | N/A |
| PQL46 | (Liu et al., 2018) |  |

## EXPERIMENTAL MODEL AND SUBJECT DETAILS

### Experimental Materials

*C. elegans* strains were cultured and maintained as described (Brenner, 1974). Details and a complete list of strains in this study are shown in Table S1. The HEK293FT cell line was obtained from ATCC (American Type Culture Collection).

## METHOD

### Molecular Biology and Transgenesis

We used standard molecular biology techniques for cloning and plasmid construction. Most of the plasmid constructs were generated in pSM vector backbone, more details see Key Resources Table. Germline transformation of *C. elegans* was performed using standard techniques (Mello and Fire, 1995). The co-injection marker plasmid and the concentration are as follows, *pCFJ104* at 5 ng/μl, *pCFJ90* at 2.5 ng/μl, *odr-1::gfp* at 60 ng/μl, or *odr-1::rfp* at 60 ng/μl.

### Scoring Migration Defects of QL Neuroblast Descendants

PQR was assayed using *lqIs58[gcy-32::cfp]* in L4 animals. PVM and AVM were visualized with *zdIs5 [Pmec-4::gfp]*. SDQL was assayed using extrachromosomal arrays *Pgcy-35::mcherry*. Migration defects in QL neuroblast descendants (PQR, PVM, SDQL) depend on neuron positions in worm. We quantified the neuron position in L4 or young adult animals. We chose never ring, vulva and anus as fiduciary markers. The relative position of PQR was calculated as the distance between never ring and PQR divided by the distance between never ring and anus. The relative positions of SDQL and PVM were calculated as the distance between SDQL or PVM and vulva divided by the distance between never ring and anus. Neurons posterior to vulva were calculated as positive values and neurons anterior to vulva are calculated as negative values respectively.

### Imaging

Detail imaging procedures were performed as described (Wang et al., 2021) with slight modifications as follows. Animals were anesthetized with 5 mmol/L levamisole in M9 buffer, and then mounted on 2% (w/v) agarose pads. PQR neuron in L4 animal was visualized with *lqIs58[gcy-32::gfp]* and imaged using OLYMPUS BX53 fluorescent microscope with a UPlanSApo 20x/0.75 objective. The CellSens software is used to process the image. To analyze the position of other neurons, *zdIs5[Pmec-4::gfp]* marked AVM and PVM in L2 animals, extrachromosomal arrays *(Pgcy-35::mcherry)* marked SDQL in L4 animals, we used a Nikon Spinning Disk confocal microscope (TI2+CSU+W1) with Photometrics Prime 95B camera, W1 spinning disk head, the 488/561 nm excitation laser and a 60×/1.40 N.A oil immersion objective. Maximum-intensity projections were generated using ImageJ (NIH).

To image the fluorescent marker strains *rdvIs1[Pegl-17::mcherry], muIs16 [mab-5::GFP], ki[gfp::catp-8], muIs49[egl-20::GFP]*, and transgenic arrays *Pegl-20::nls::mcherry*, we used Zeiss Axio Observer Z1 microscope (Carl Zeiss) equipped with an alpha Plan-Apochromat 63x/1.46 NA objective. ImageJ software (NIH) was used to process the images.

### Time-Lapse Imaging of Neuronal Migration

*C. elegans* L1 larvae were anesthetized with 2.5 mmol/L levamisole in M9 buffer, and then mounted on 2% (w/v) agarose pads. Slides were sealed with 2:1 vaseline/paraplast tissue embedding medium. Images were acquired using a Zeiss Axio Observer Z1 microscope (Carl Zeiss) equipped with an alpha Plan-Apochromat 63x/1.46 NA objective, Yokogawa camera adapter, 561 nm laser and Hamamatsu camera. QL neuroblast migration was visualized using *rdvIs1[Pegl-17::mCherry]*, a stack at 1 um intervals with mCherry exposure time of 3.5 s at every 15 min. ImageJ software (NIH) was used to process the images.

### CRISPR/Cas9-Mediated Genomic Editing

To generate *yan125* mutant by CRISPR/Cas9 as described (Dickinson et al., 2013), *Peft-3::cas9* (50 ng/μl), *U6::egl-20-sg#1* (25 ng/μl, target sequence:5’-TATTTGTTCTCCTCGTTTA-3’), *U6::egl-20-sg#2* (25 ng/μl, target sequence:5’-AAACTTACAGCCAGTTATA-3’) and *pCFJ104* (5 ng/μl) were injected into N2 strain. F1 worms were screened for successful knock-out by PCR. F2 worms were cloned out without co-injection marker, genotyped by PCR, and sequenced to confirm.

To replace EGL-20 (1-18 aa) signal sequence with PAT-3 (1-20 aa) signal sequence, coding sequence of PAT-3 signal sequence was inserted in *pSM-egl-20* vector via long primers. Two homologous arms were 679 bp (5’-arm) and 654 bp (3’-arm) respectively. Repair template (50 ng/μl), *U6::egl-20-sg#1* (25 ng/μl) *U6::egl-20-sg#2* (25 ng/μl), pCFJ104 (5 ng/μl) were injected into *catp-8(yan22)*. F1 worms were screened for successfully insertion using PCR, F2 worms without co-injection marker were cloned out, genotyped by PCR, and verified by sequencing.

To rescue the PQR migration defects in *catp-8(yan22)*, a single copy insertion of *yanTi9[Pcatp-8::catp-8]* was generated by CRISPR-Cas9 in *catp-8(yan22)*. The promoter of *catp-8* from N2 genomic DNA and *catp-8* cDNA were amplified and inserted in *ttTi5605* vector. Repair template (50 ng/μl), *pDD122* (50 ng/μl), *pCFJ104* (5 ng/μl) and *pCFJ90* (2.5 ng/μl) were injected into *catp-8(yan22)*. F1 worms were screened for successfully insertion using PCR. F2 worms without co-injection marker were cloned out, genotyped by PCR and confirmed by sequencing.

### Western Blot

We synchronized early L1 larvae by hatching them in M9 buffer, and resuming feeding on regular OP50 NGM plates till they grew to L4 stage. Worms were harvested and washed for three times with M9 buffer, and the pellet was lysed in Laemmli sample buffer (32.9 mM Tris-HCl, pH 6.8, 13.2% glycerol, 1.05% SDS, 2.5% 2-mercaptoethanol and 0.005% bromophenol blue). Pellets were repeatedly boiled for 5 min and then vortexed for 1 min, until most of the worms were broken. The lysates were spun at 13,000 rpm for 10 min at 4°C. Then supernatants were collected and analyzed using western blots with mouse monoclonal anti-GFP (1:1000, Abmart), Anti-β-Actin pAb-HRP-DirecT (1:5000, Medical & Biological Laboratories) and HRP-conjugated Rabbit antibody to mouse (1:10,000, Sigma).

Plasmids were transfected into control or ATP13A1 KO cells, and then cell lysates were extracted by NP40 lysis buffer (150 mM NaCl, 0.5% NP40 and 50 mM Tris-HCl pH 7.4, 1mM EDTA, 10% glycerol) containing 1% Protease Inhibitor stock (Bimake) for 30 min on ice. The soluble fraction of the cell lysates was isolated via centrifugation at 12,000 rpm for 10 min at 4°C. Laemmli sample buffer was added to the supernatant. Then the samples were heated at 65°C for 10 min, resolved via SDS-PAGE gel electrophoreses, and analyzed using Western blot with the rabbit polyclonal anti-ATP13A1 (1:2000, proteintech), mouse anti-FLAG (1:1000, sigma), HRP-conjugated donkey anti-rabbit (1:10000, Jackson Immunoresearch), and HRP-conjugated Rabbit antibody to mouse (1:30,000, Sigma).

### WNT1 secretion assay in HEK293FT cells

WNT1 secretion assay were performed as described (Torpe et al., 2019). Control and *ATP13A1* KO cells were cultured overnight in DMEM containing serum and L-glutamine. Cells were transfected with plasmids encoding WNT1::FLAG::Opsin as indicated in Figure 4A. 24 h after transfection, the media was replaced with 500ul of serum-free DMEM. Cells were incubated at 37°C for 16 h before cells and media were collected. The cells were lysed with NP40 lysis buffer (150 mM NaCl, 0.5% NP40 and 50 mM Tris-HCl pH 7.4, 1 mM EDTA, 10% glycerol). The media were collected performed with anti-FLAG beads (Sigma). Samples were eluted off from the beads via FLAG peptide (Sigma). The lysate was treated as above.

### Real-Time RT-PCR

We synchronized L1 larvae by bleaching adult animals to obtain eggs to hatch on NGM plate with OP50. L1 worm was collected and lysed in 1 mL Trizol reagent (Vazyme) by freeze and thaw. Total RNA extraction was followed by reverse-transcription with random primers and HiScript II Q RT SuperMix for qPCR (Vazyme). Relative abundance of *egl-20* and actin was quantified with real-time PCR using TB Green Premix Ex TaqII (Tli RNase H Plus, Takara).

### Statistical Analysis

Quantification of neuron (PQR, PVM, SDQL) positions along the anterior-posterior axis of the worm body was One-way ANOVA followed by Tukey’s multiple comparisons test (Prism; GraphPad Software). For western blot, ImageJ Software (NIH) was used to analyze the grayscale values from three biological repeats. Statistical comparisons were conducted using Tukey’s multiple comparisons test. For data in Figure 3B and Figure 3H, Student’s t test was used. Statistical significance is indicated as n.s., not significant; *, *P* < 0.05; **, *P* < 0.01; ***, *P* < 0.001; ****, *P* < 0.0001.

